## Supplement 1 for "Human hypertrophic cardiomyopathy mutation R712L suppresses the working stroke of cardiac myosin and can be rescued by omecamtiv mecarbil"

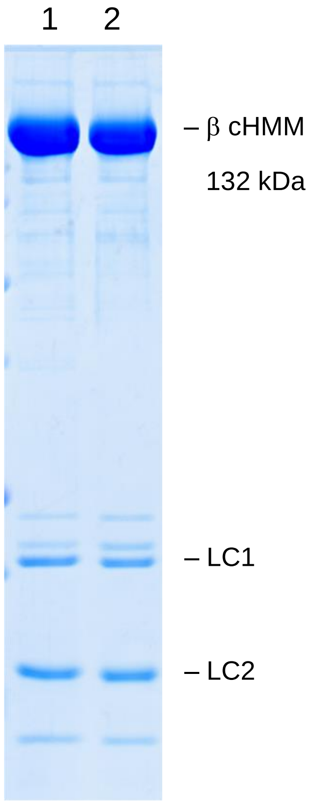


**Figure 1—Supplement 1.** Purified wild type human β-cardiac HMM (WT-myosin) and the R712L HCM variant (R712L-myosin) are routinely analyzed by SDS PAGE. Lane 1, WT-myosin; Lane 2, R712L-myosin. The purified proteins consist of a 132 kDa heavy chain and associated myosin light chains LC1 and LC2.

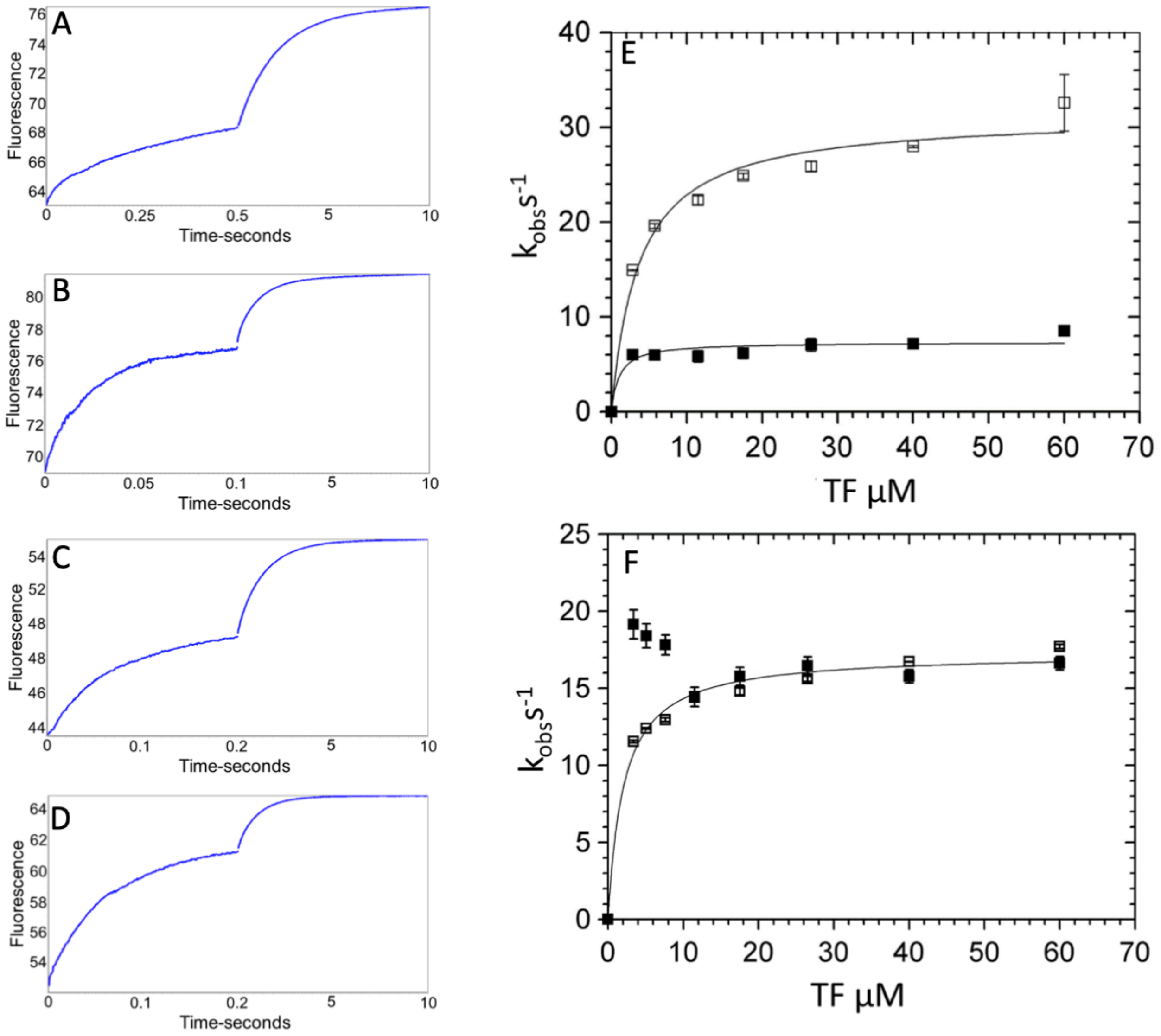


**Table 1—Supplement 1. Kinetics of phosphate dissociation from thin filament (TF)-activated β-cardiac myosin.** Thin filament activation of phosphate dissociation was measured via the change in MDCC-PBP fluorescence (see Methods). Briefly, 3 µM β-cardiac myosin was first mixed with 2 µM ATP, held in a delay line for 2 s, and then mixed with thin filaments to accelerate $P_i$ release. Final concentrations in the flow cell were 0.75 µM myosin, 0.5 µM ATP, 0-60 µM actin in native TFs, 5 mM MOPS (pH 7.2), 2 mM $MgCl_2$, 10 mM KCl, 0.05 mM $CaCl_2$, 0.25% DMSO, 0 or 50 µM OM, 5 µM MDCC-PBP, 0.1 mM 7-methylguanosine, and 0.01 unit/mL purine nucleoside phosphorylase at 20°C. (**A-D**) Representative traces showing the change in MDCC-PBP fluorescence with 60 µM actin in TFs in the presence and absence of OM for WT- and R712L-myosin. The data were fit to the sum of two-exponentials for: **(A)** WT-myosin (7.8 $s^{-1}$, 0.52 $s^{-1}$), **(B)** WT-myosin with OM (42 $s^{-1}$, 1.0 $s^{-1}$), **(C)** R712L-myosin (17 $s^{-1}$, 0.78 $s^{-1}$), and **(D)** R712L-myosin with OM (17 $s^{-1}$, 1.0 $s^{-1}$). **(E-F)** Kinetics of phosphate release for WT **(E)** and for R712L-myosin **(F)** as a function of actin subunit concentration in porcine native thin filaments. (■) - without OM; (□) - with OM. Solid lines are hyperbolic fits with constants reported in Table 1.

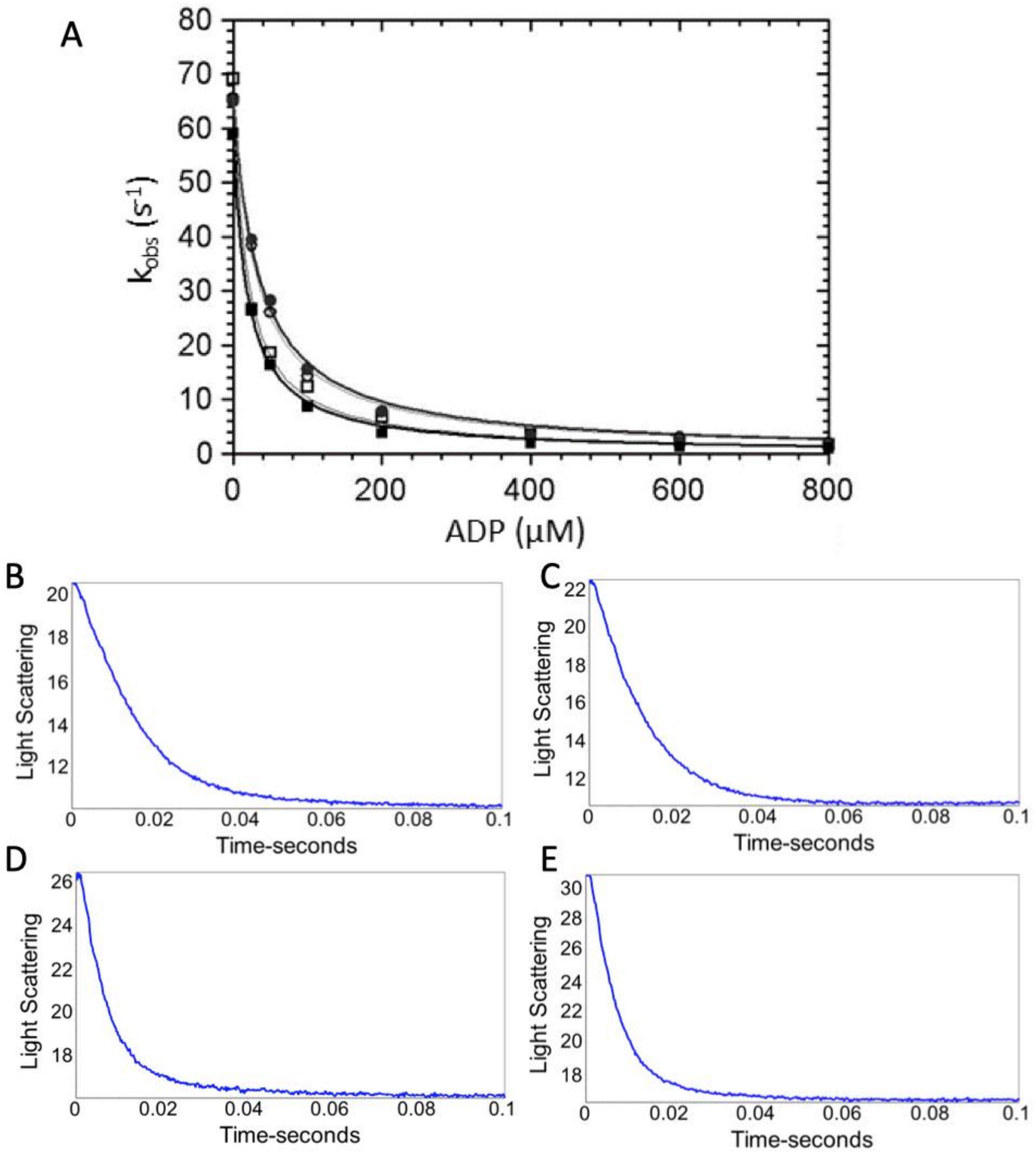


**Table 1—Supplement 2. ADP Dissociation from actomyosin. (A)** ADP dissociation from actomyosin was measured via light scattering (see Methods). Briefly, stopped flow was used to quickly mix actomyosin and ADP with ATP (30 µM final concentration), and a decrease in light scattering is due to dissociation of actomyosin is known to be limited by ADP dissociation in WT myosin. Final concentrations of 1.25 µM myosin, 3 µM actin, 0 µM – 800 µM ADP, 30 µM ATP, 2 mM $MgCl_2$, 25 mM KCl, 0.25% DMSO and either 0 or 50 µM OM in 5 mM MOPS (pH 7.2). (■) WT-myosin, (□) WT myosin + 50 µM OM, (●) R712L-myosin, (○) R712L-myosin + 50 µM OM. **(B-E)** Maximum rates of ADP release were obtained with final concentrations in the cell of 1.25 µM myosin, 3 µM actin, 140 µM ADP, 2 mM ATP, 5 mM MOPS (pH 7.2), 2 mM $MgCl_2$, 25 mM KCl, 0.25% DMSO and either 0 or 50 µM OM. Representative traces from: **(B)** WT-myosin ($k_{obs}$ = 74 $s^{-1}$), **(C)** WT-myosin + 50 µM OM ($k_{obs}$ = 85 $s^{-1}$), **(D)** R712L-myosin ($k_{obs}$ = 141 $s^{-1}$), and **(E)** R712L-myosin + 50 µM OM ($k_{obs}$ = 147 $s^{-1}$). Average values of $k_{-AD}$ from multiple experiments are reported in Table 1.

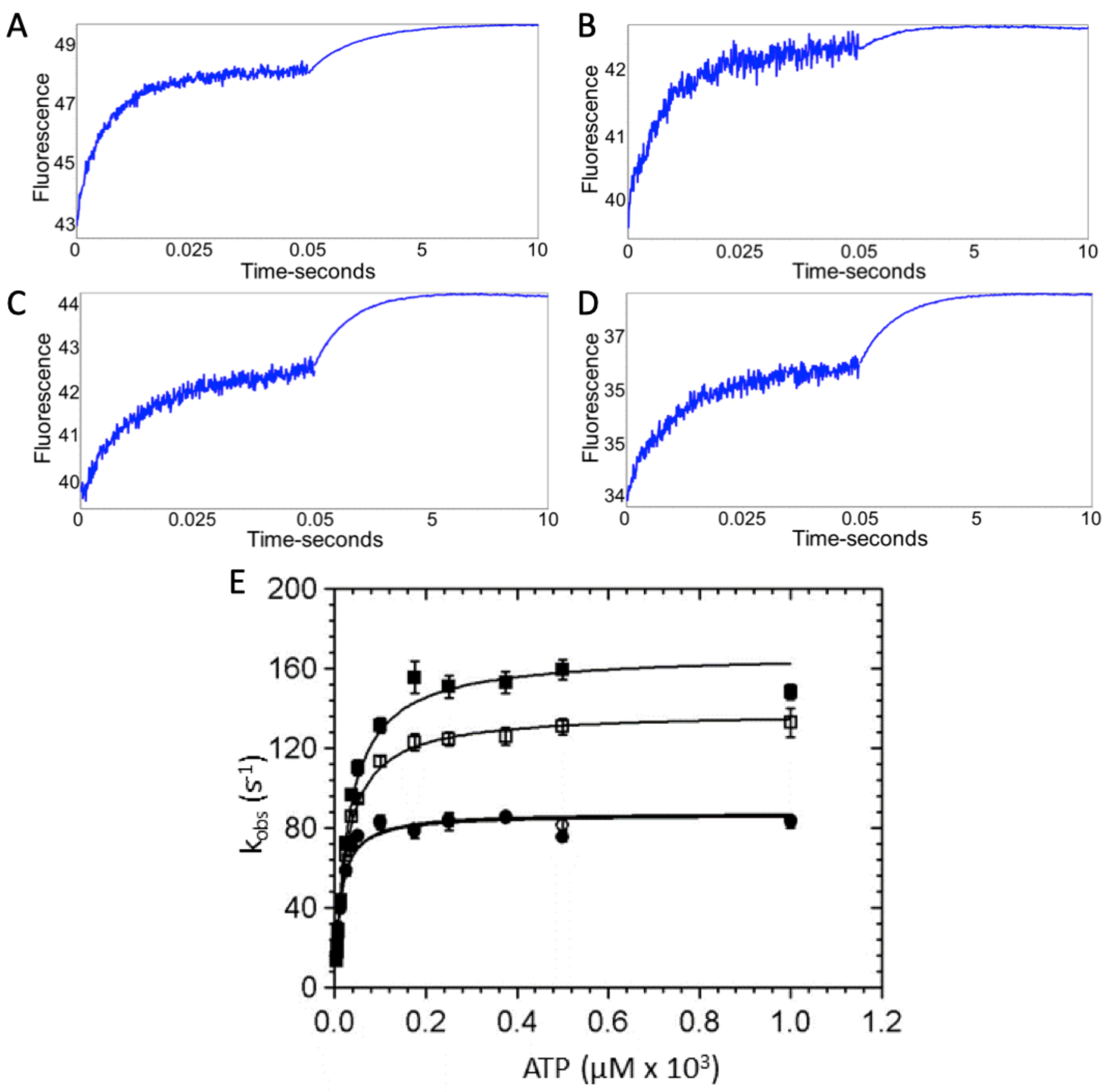


**Table 1—Supplement 3. ATP binding to β-cardiac myosin measured by intrinsic tryptophan fluorescence.** Myosin was mixed with increasing concentrations of ATP via stopped flow and intrinsic tryptophan fluorescence was monitored. Upon ATP binding intrinsic tryptophan fluorescence of the myosin increases. **(A-D)** Representative traces showing the change in fluorescence with 175 µM ATP in the presence and absence of OM for WT- and R712L-myosin are shown and are best fit by the sum of two exponentials: **(A)** WT-myosin (139 $s^{-1}$, 0.48 $s^{-1}$), **(B)** WT-myosin with OM (115 $s^{-1}$, 0.82 $s^{-1}$) **(C)** R712L-myosin (80 $s^{-1}$, 0.77 $s^{-1}$), **(D)** R712L-myosin with OM (84 $s^{-1}$, 0.70 $s^{-1}$). **(E)** Rate of ATP binding to myosin as a function of ATP concentration. (■) WT-myosin, (□) WT-myosin + 50 µM OM, (●) R712L-myosin, (○) R712L-myosin + 50 µM OM. Final concentrations in the cell were 1.25 µM myosin, 3.75 µM to 1 mM ATP, 5 mM MOPS (pH 7.2), 2 mM $MgCl_2$, 25 mM KCl, 0.25% DMSO, and 0 or 50 µM OM. Reactions were carried out at 20 °C. Solid lines are hyperbolic fits, with constants reported in Table 1.

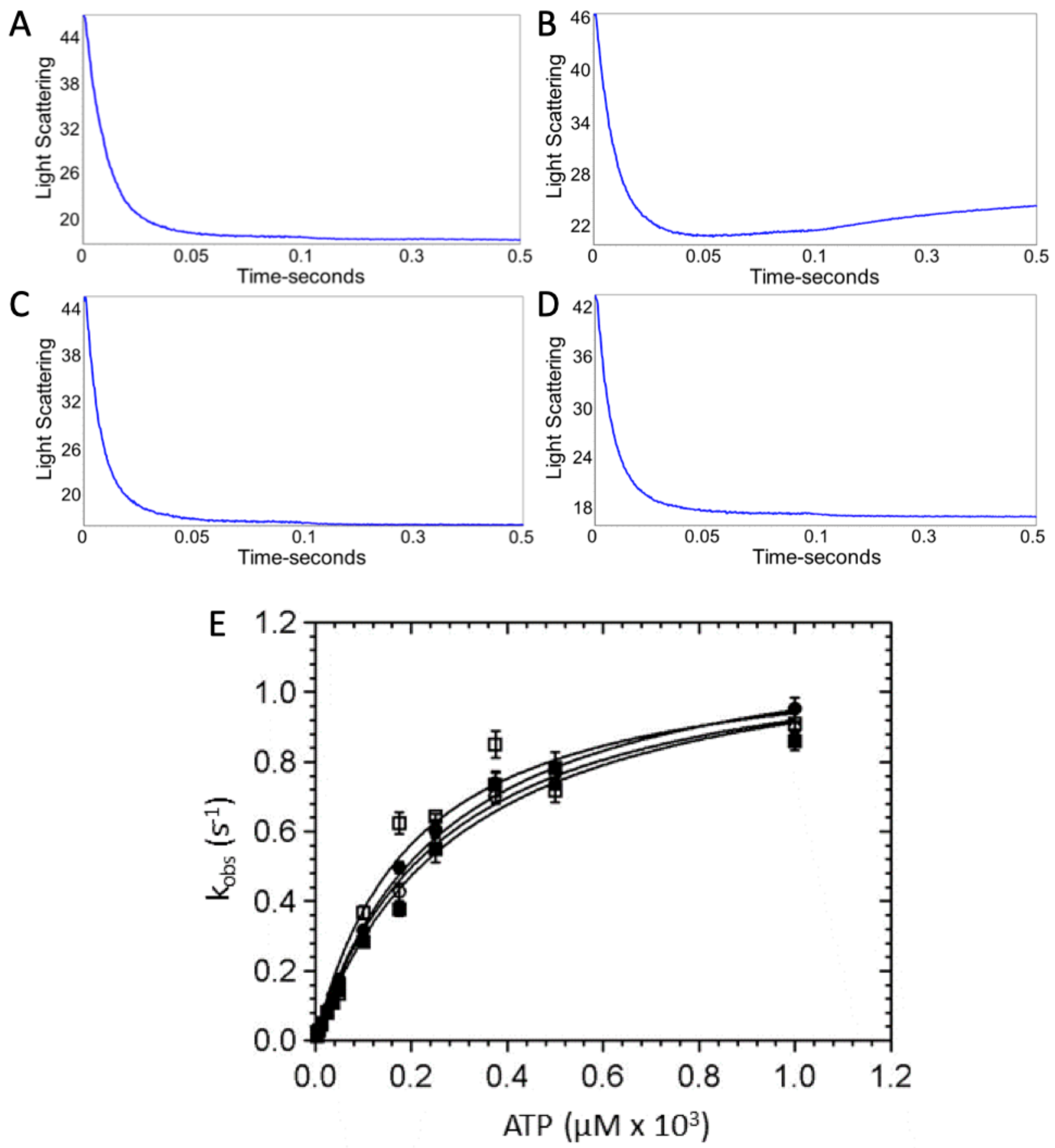


**Table 1—Supplement 4. ATP-induced dissociation of β-cardiac myosin from actin**. A decrease in light scattering due to actomyosin dissociation was observed upon mixing 2.5 μM myosin and 6 μM actin with increasing concentrations of ATP in a stopped flow device. Representative traces showing the change in light scattering in arbitrary units at 50 μM ATP in the presence and absence of OM for WT- and R712L-myosin. Representative traces for: **(A)** WT-myosin (105 $s^{-1}$, 19 $s^{-1}$), **(B)** WT-myosin + OM (108 $s^{-1}$, 2.4 $s^{-1}$), **(C)** R712L-myosin (155 $s^{-1}$, 32 $s^{-1}$), and **(D)** R712L-myosin + OM (147 $s^{-1}$, 29 $s^{-1}$). **(E)** Rate of actomyosin dissociation as a function of ATP concentration. (■) WT-myosin, (□) WT-myosin + 50 μM OM, (●) R712L-myosin (○) R712L-myosin + 50 μM OM. Solid lines are fitted hyperbolae with constants given in Table 1.. Experimental conditions in the cuvette: 1.25 μM myosin, 3 μM actin, 3.75 μM - 1 mM ATP, 5 mM MOPS (pH 7.2), 2 mM $MgCl_2$, 25 mM KCl, 0.25% DMSO and either 0 or 50 μM OM, 20 °C.

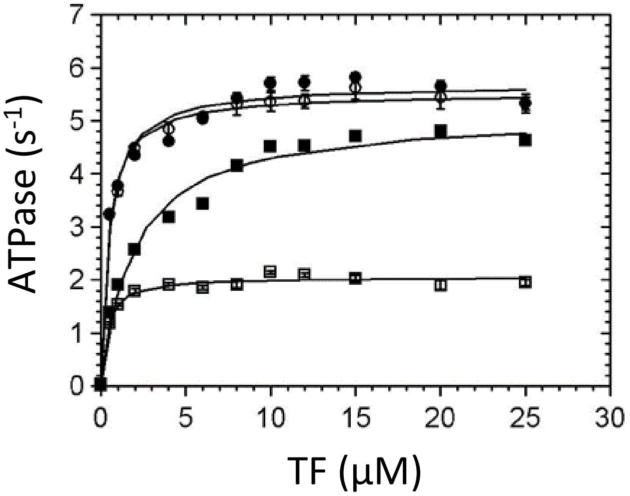


**Table 1—Supplement 5. Thin-filament activation of steady state ATPase activity.** Steady-state ATPase rates were measured at using an NADH-coupled assay at 25 °C (5 mM MOPS pH 7.2, 2 mM $MgCl_2$, 1 mM DTT, 2 mM ATP, 0.01 mM EGTA, 0.1 mM $CaCl_2$, 0.5% DMSO and either 0 or 50 μM OM). 0.02-0.025 μM myosin and 0-25 μM actin concentration in thin filaments (TFs) were used. (■) WT myosin, (□) WT myosin + 50 μM OM, (●) R712L, (○) R712L + 50 μM OM. Solids lines are fits of the data by hyperbolic functions, with constants given in Table 1.

**Movie Legends:**

**Movie 1:** The four panels of this movie illustrate the contrasting effect of OM on WT- and R712L-myosin. All panels are from the same experiment with the same surface density of cardiac HMM and correspond to 100 frames captured at 5 frames/s and played back in the short clip at 30 frames/s. The faster playback frame rate is necessary to show the slow movement of the WT-myosin at 12.5 µM OM. Actin filament velocities measured for each condition are: WT-No OM: 1.45 µm/s, R712L-No OM: 0.27 µm/s, WT+12.5 µM: OM 0.06 µm/s, and R712L+50 µM OM: 0.81 µm/s. The last image in each panel is a maximum projection of the image stacks revealing the tracks followed by the individual filaments. Each panel is 96 µm x 73 µm.

**Movie 2:** The R712L mutation affects the coupling of the motor domain relay helix to the converter/lever arm. Animated MD trajectory showing 100 ns equilibration of β-cardiac myosin (PDB: 6FSA) with (blue) and without (grey) the R712L mutation. The initial structure is shown in red, and the cylinders show the positions of the lever-arm helix axes. After 100 ns, the lever-arm helix is pulled for 125 ns (see Methods). The arrow shows the force vector, as pulling occurred toward the pointed-end and parallel to the long-axis of a hypothetical bound actin filament.

**Movie 3:** Morph created in PyMOL showing interpolated trajectory between the initial structure (PDB: 6FSA) and the last frame of the 100 ns equilibration (Figure 2, Movie 2). The position of OM (yellow) was determined from PDB: 4PA0. The converter domain and lever-arm helix are shown in dark blue. A clash is shown that would occur between the OM and amino acid residues P710-I713 of the converter if OM remained in its WT binding site.
